# Three new species in the Onygenaceae isolated from marine environment

**DOI:** 10.1101/2023.12.31.573767

**Authors:** Meng-Meng Wang, Qi Li, Wei Li, Lei Cai

## Abstract

Several fungal isolates affiliated with *Aphanoascus* and *Chrysosporium* were obtained from the sediments of Chinese seas by us. Among them, one new *Aphanoascus* species (*A. sedimenticola*) and two new *Chrysosporium* species (*C. microsporum, C. sphaerospermum*) were taxonomically identified and recognized based on phylogenetic analyses and morphological characteristics. This paper provided the detailed descriptions and illustrations of the three new taxa.

## Introduction

The genus *Aphanoascus* Zukal was introduced in 1890, with *A. fulvescens* as type species (Zukal 1890; Apinis 1968). The nomenclature of *Aphanoascus* has been changed many times historically. Cooke published *Badhamia fulvescens* in 1875 and renamed as *Eurotium fulvescens* in 1879 based on description of *E. stercorarium* Hansen (Cooke 1875; Cooke and Ellis 1879; Hansen 1877). Zukal (1890) recognized *E. stercorarium* differs from other *Eurotium* species and introduced the genus *Aphanoascus* with the only species *A. cinnabarinus*. Hansen re-examined type specimen of *E. stercorarium* and proposed a new genus *Anixiopsis* with *An. sterocoraria* as type species (Hansen 1897). De Vries (1964) considered *A. cinnabarinus* as a synonym of *An. stercoraria*. However, Apinis (1968) suggested that the above two species can be identical in morphology, and proposed a new combination *A. fulvescens* to replace *A. cinnabarinus* as type species of *Aphanoascus*. Currently, there are 18 species accepted in this genus (Wijayawardene et al. 2022).

The genus *Chrysosporium* was introduced by Corda (1833), with *C. merdarium* as the type species (Oorschot 1980). Historically, taxonomic boundary and description of this genus has been changed several times. Saccardo (1901) regarded this genus as a synonym of *Sporotrichum* Link ex Fr. Carmichael (1962) redefined *Chrysosporium* and regarded four genera *Myceliophthora* Costantin, *Geomyces* Traaen, *Blastomyces* Gilchrist & W.R. Stokes and *Emmonisia* Cif. & Montemart. as its synonyms. Oorschot (1980) re-evaluated *Chrysosporium* and confirmed 22 species based on morphological characteristics. With the application of molecular systematics in the taxonomy of *Chrysosporium* in the past decade, more new species have been reported based on morphology and ITS phylogeny (Han et al. 2013; Zhang et al. 2017; Li et al. 2019; Zhao et al. 2018; Zhang et al. 2020). The genus currently comprises 67 species (Wijayawardene et al. 2022 Zhang et al. 2020).

Members in the family Onygenaceae, including *Aphanoascus* and *Chrysosporium*, are widely distributed and exists in various habitats such as air, sludge, waste water and animal faeces (Deshmukh 1999, Zhang et al. 2016, Zhang et al. 2020), especially in rich keratin substance (Zhang et al. 2017). However, their distribution and species diversity in marine environments are poorly known so far, except one species *C. merdarium* reported (Jones et al. 2019). During our investigation of fungal diversity from Chinese marginal seas, several fungal isolates identified as *Aphanoascus* and *Chrysosporium* were obtained from intertidal and coastal sediments. Here, we introduced them as three new species with the evidences of morphological characteristics and phylogenetic analyses.

## Materials and Methods

### Sample collection and fungal isolation

Sediment samples were collected from intertidal zones of Fujian, Guangdong and Shandong provinces following protocol of Wang et al. (2024), and from coastal regions of Bohai Sea following protocol of Wang et al. (2023ME), respectively. Fungi were isolated from sediment samples using the dilution plate method and the direct isolation method following protocol of Wang et al. (2024).

All isolates examined in this study were deposited in Wei Li’s personal culture collection (WL). Type specimens of new species were deposited in the Fungarium of the Institute of Microbiology (HMAS), with the ex-type living cultures in the China General Microbiological Culture Collection Center (CGMCC).

### Morphological observation

The isolates studied were incubated on Potato Dextrose Agar (PDA; each 1 L medium containing potato 200.0 g, dextrose 20.0 g, agar 20.0 g, and seawater 1 L) and Czapek-Dox Agar (CDA; each 1 L medium containing sodium nitrate 3.0g, dipotassium phosphate 1.0g, magnesium sulfate 0.5g, potassium chloride 0.5g, ferrous sulfate 0.01g, dextrose 30.0 g, agar 20.0g, and seawater 1L). After 7 days of incubation in the dark, culture characteristics including colony morphology, pigmentation, and odour were observed. Colours were assessed according to the colour charts of Kornerup and Wanscher (1978). Micromorphological characteristics were examined and photo-documented using water as a mounting medium under an Olympus BX53 microscope with differential interference contrast (DIC) optics. For each species, 30 conidiophores, 30 conidiogenous cells and 50 conidia were mounted and measured randomly.

### DNA extraction and amplification

Genomic DNA was extracted from fungal mycelia grown on PDA, using a modified CTAB protocol as described in Guo et al. (2000). The internal transcribed spacer region (ITS) was amplified using primer pairs ITS5/ITS4 (White et al. 1990). Amplification reactions were performed following protocol described in Wang et al. (2024). The PCR products were visualised on 1% agarose electrophoresis gel. Sequencing was performed bidirectionally and conducted by the BGI Write Company (Beijing, China). Consensus sequences were obtained using SeqMan of the Lasergene software package v. 14.1 (DNAstar, Madison, WI, USA).

### Phylogenetic analyses

The sequences of the *Aphanoascus* and *Chrysosporium* strains examined in this study and the references strains are listed in Table 1. The ITS sequences were aligned using MAFFT v. 7(Katoh et al. 2018), and the alignments were manually adjusted where necessary. The best-fitting nucleotide-substitution models according to the Akaike Information Criterion (AIC) were selected using jModelTest v. 2.1.7 (Posada et al. 2008; Darriba et al. 2012). Alignment derived from this study were deposited in TreeBASE (submission ID xxxxx), and taxonomic novelties were deposited in FungalNames.

Phylogenetic analyses of the ITS dataset were performed using Bayesian inference (BI) and maximum-likelihood (ML) methods. The BI analysis was conducted using MrBayes v. 3.2.1 (Huelsenbeck et al. 2001) following the protocol of Wang et al. (2019), with optimization of each locus treated as a partition in combined analyses, based on the Markov Chain Monte Carlo (MCMC) approach (Ronquist et al. 2012). All characters were equally weighted, and gaps were treated as missing data. The stationarity of the analyses was determined by examining the standard deviation of split frequencies (<0.01) and –ln likelihood plots in AWTY (Nylander et al. 2008). The ML analyses were conducted using PhyML v. 3.0 (Guindon et al. 2010), with 1000 bootstrap replicates. The general time reversible model was applied with an invariable gamma-distributed rate variation (GTR+I+G).

## Results

### Phylogenetic analyses

Analyses of the *Aphanoascus* and *Chrysosporium* phylogeny were conducted by using the ITS dataset. For the BI and ML analyses, the GTR+I+G model was selected. The phylogeny showed that two of our isolates (WL00820 and WL04860) and a reference isolate (NBRC31723) formed a single clade in the genus *Aphanoascus*, while three isolates (WL02687, WL02759 and WL02839) and two isolates (WL03114, WL03341) formed two distinct clades in the genus *Chrysosporium* (Figure 1).

**Figure 1.**
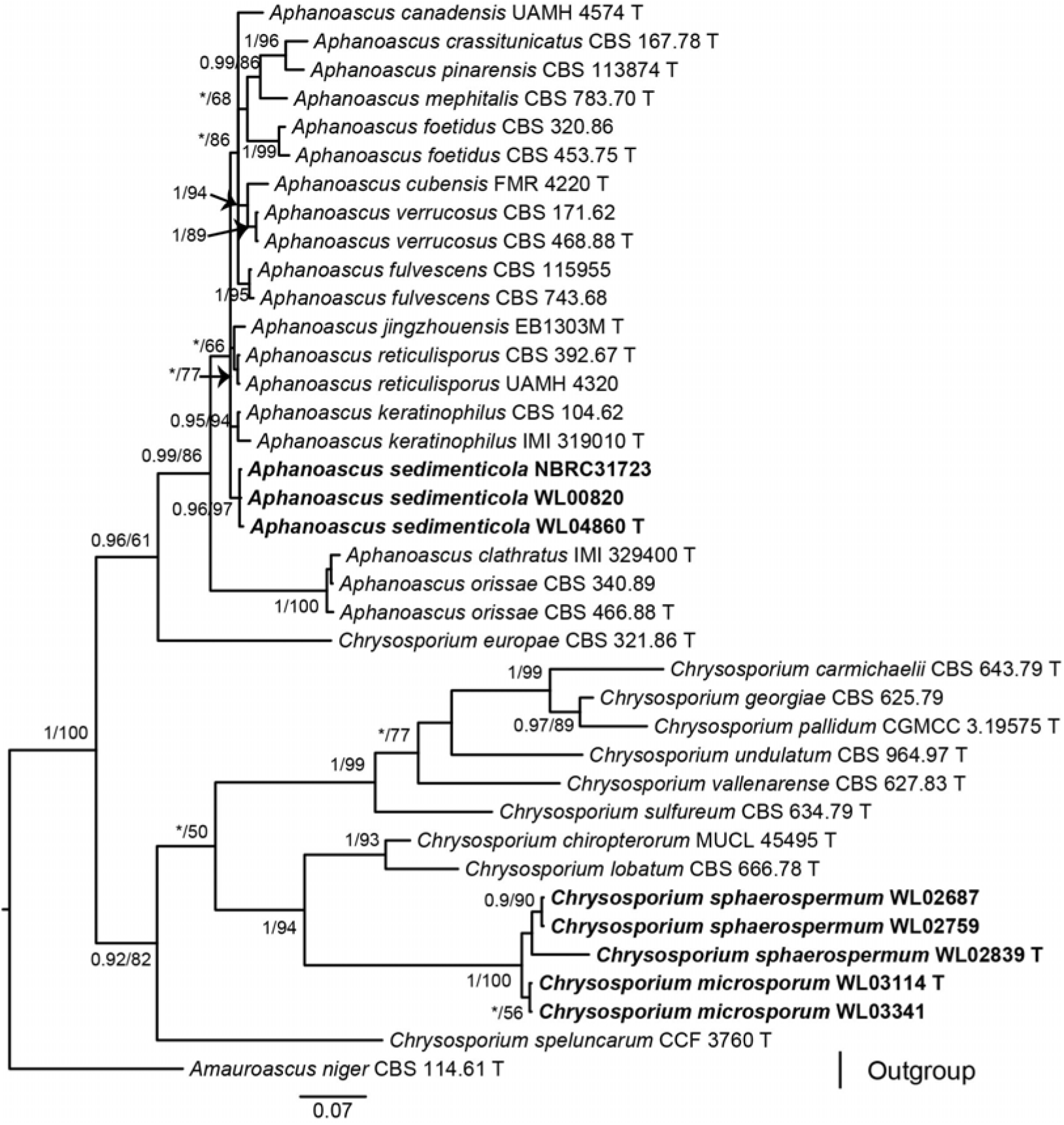
Fifty percent majority rule consensus tree from a Bayesian analysis based on ITS sequences showing the phylogenetic relationships of *Aphanoascus* and *Chrysosporium*. The Bayesian posterior probabilities (PP > 0.9) and PhyML bootstrap support values (BS > 50%) are displayed at the nodes (PP/BS). The tree was rooted to *Amauroascus niger* (CBS 114.61 T). Ex-type cultures are indicated with “T”. New species introduced in this study are marked in bold.

Descriptions and illustrations of new species

*Aphanoascus* Zukal, Ber. dt. bot. Ges. 8: 295 (1890)

*Aphanoascus sedimenticola* M.M. Wang, W. Li & L. Cai sp. nov. Fig. 2

**Figure 2.**
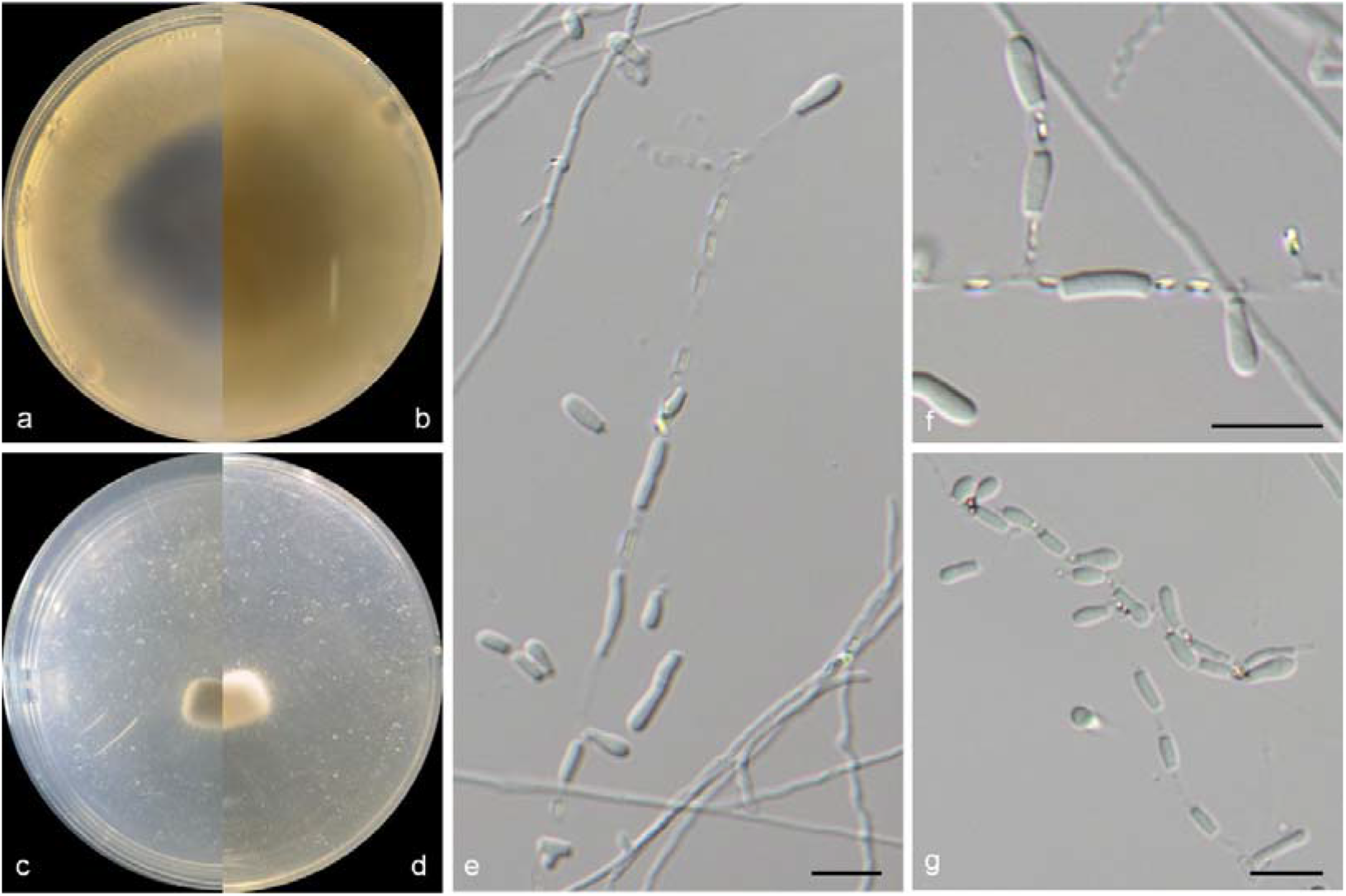
morphological photograph of *Aphanoascus sedimenticola* (ex-type OUCMB□_19_1607). a-d: colonies on PDA and CDA after 7d; e-f: Conidiophores and conidiogenous cells; g: Conidia. Bars: e-g = 10 μm.

FungalNames: IF XXXXXX.

*Etymology*: named after the habitat of the type specimen of this species, sediment.

*Typus*: CHINA, Bohai Sea, from sediment, Aug. 2019, Y.Y. Ma (holotype designated here, dried culture on SNA; culture ex-type CGMCCXXXX = WL04860).

Ascomata not observed. Peridial hyphae difficult to distinguished from aerial hyphae, septate, branched and anastomosed, terminated by short blunt prominences, smooth, thick-walled, hyaline, 3.5–5 µm diam. Arthroconidia abundant, intercalary, laterial or terminal, unicellular, hyaline; intercalary conidia cylindrical or ellipsoidal with truncated base, 4–8 × 2–3 µm; laterial or terminal conidia arised from aerial hyphae directly, racquet or clavate with truncated base, 5–7 × 2–4 µm.

Culture characteristics — Colonies on PDA 35–40 mm diam in 7 d at 25°C, pulvinate, erose, greyish purple near the centre, with white to pale grey regular margin; reverse greyish purple near the centre, white to pale grey regular margin. Colonies on CDA 25–30 mm diam in 7 d at 25°C, pulvinate, grey in the centre, with white regular margin; reverse dark grey near the centre, white near the margin.

Other examined isolates: CHINA, Bohai Sea, from sediment, Jun. 2011, C.H. Feng, X.X. Ji and Z.Z. Wang (WL00820).

*Chrysosporium* Corda, in Sturm, Deutschl. Fl., 3 Abt. (Pilze Deutschl.) 3(13): 85 (1833)

*Chrysosporium microsporum* M.M. Wang, W. Li & L. Cai sp. nov. Fig. 3 FungalNames: IF XXXXXX.

**Figure 3.**
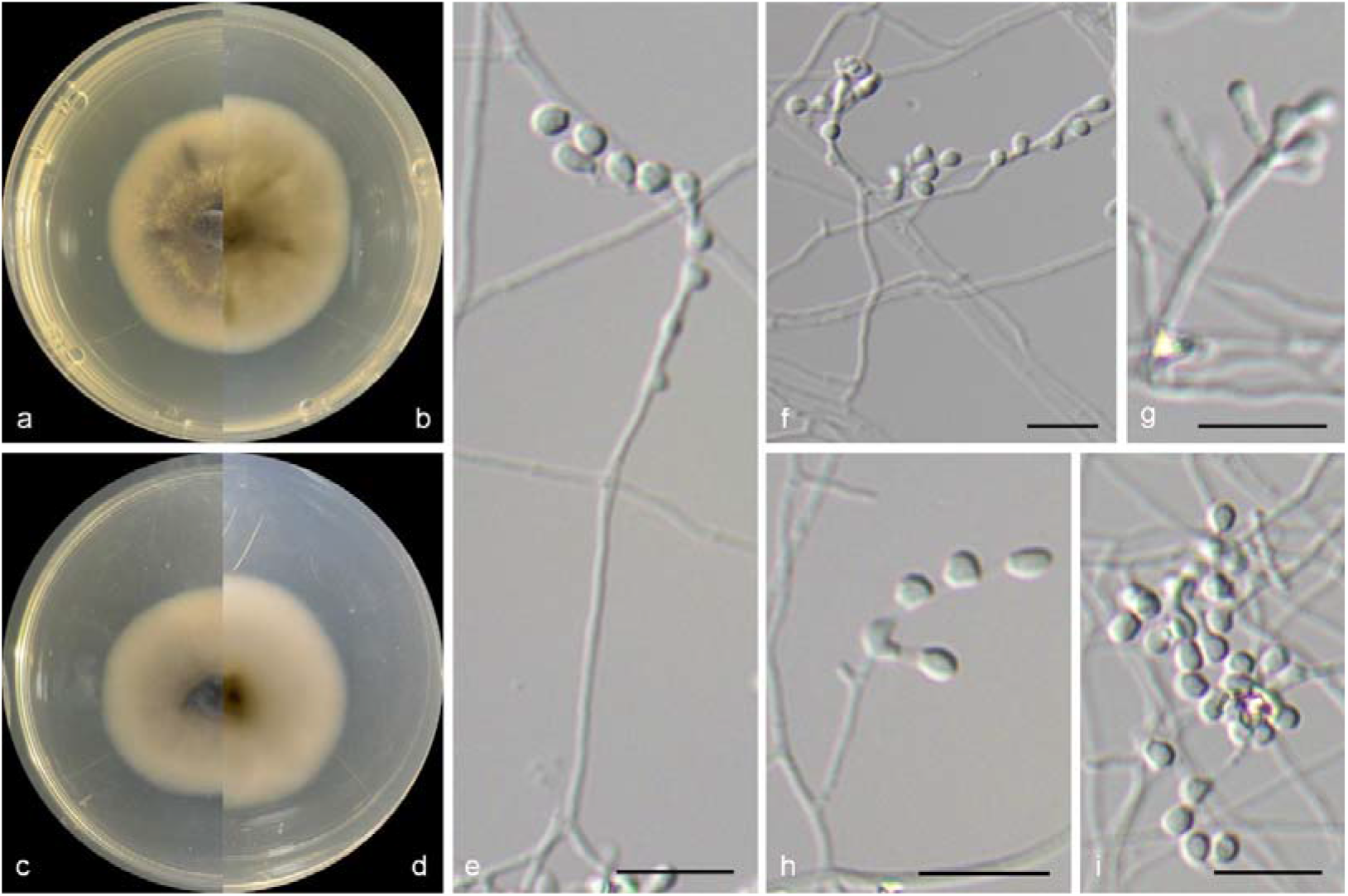
morphological photograph of *Chrysosporium microsporum* (ex-type OUCMB□_14_1201). a-d: colonies on PDA and CDA after 7d; e-h: Conidiophores and conidiogenous cells; i: Conidia. Bars: e-i = 10 μm.

*Etymology*: named after the smaller size of this species.

*Typus*: CHINA, Shandong Province, Yantai city (121°38′29.64″E 37°28′35.34″N), from sediment, Jun. 2014, X.M. Bian (holotype designated here, dried culture on SNA; culture ex-type CGMCCXXXX = WL03114).

Vegetative hyphae hyaline, septate, branched, smooth. Ascomata not observed. Peridial hyphae difficult to distinguished from aerial hyphae, septate, branched and anastomosed, terminated by short blunt prominences, smooth, thick-walled, hyaline, 2–3.5 µm diam. Arthroconidia abundant, intercalary, laterial or terminal, unicellular, hyaline; intercalary conidia cylindrical or ellipsoidal with truncated base, 3–5 × 2.5–3 µm; laterial or terminal conidia arised from aerial hyphae directly, pyriform or clavate with truncated base, 3–5 × 2.5–3 µm.

Culture characteristics — Colonies on PDA 35–40 mm diam in 7 d at 25°C, flat, greyish purple near the centre, with white to pale grey regular margin; reverse grey near the centre, white near the margin. Colonies on CDA 35–40 mm diam in 7 d at 25°C, flat, grey in the centre, with white regular margin; reverse grey near the centre, white near the margin.

Other examined isolates: CHINA, Shandong Province, Qingdao city (20°41′20.33″E 36°23′34.13″N), from sediment, Jun. 2014, X.M. Bian and W. Li (WL03341).

*Chrysosporium sphaerospermum* M.M. Wang, W. Li & L. Cai sp. nov. Fig. 4 FungalNames: IF XXXXXX.

**Figure 4.**
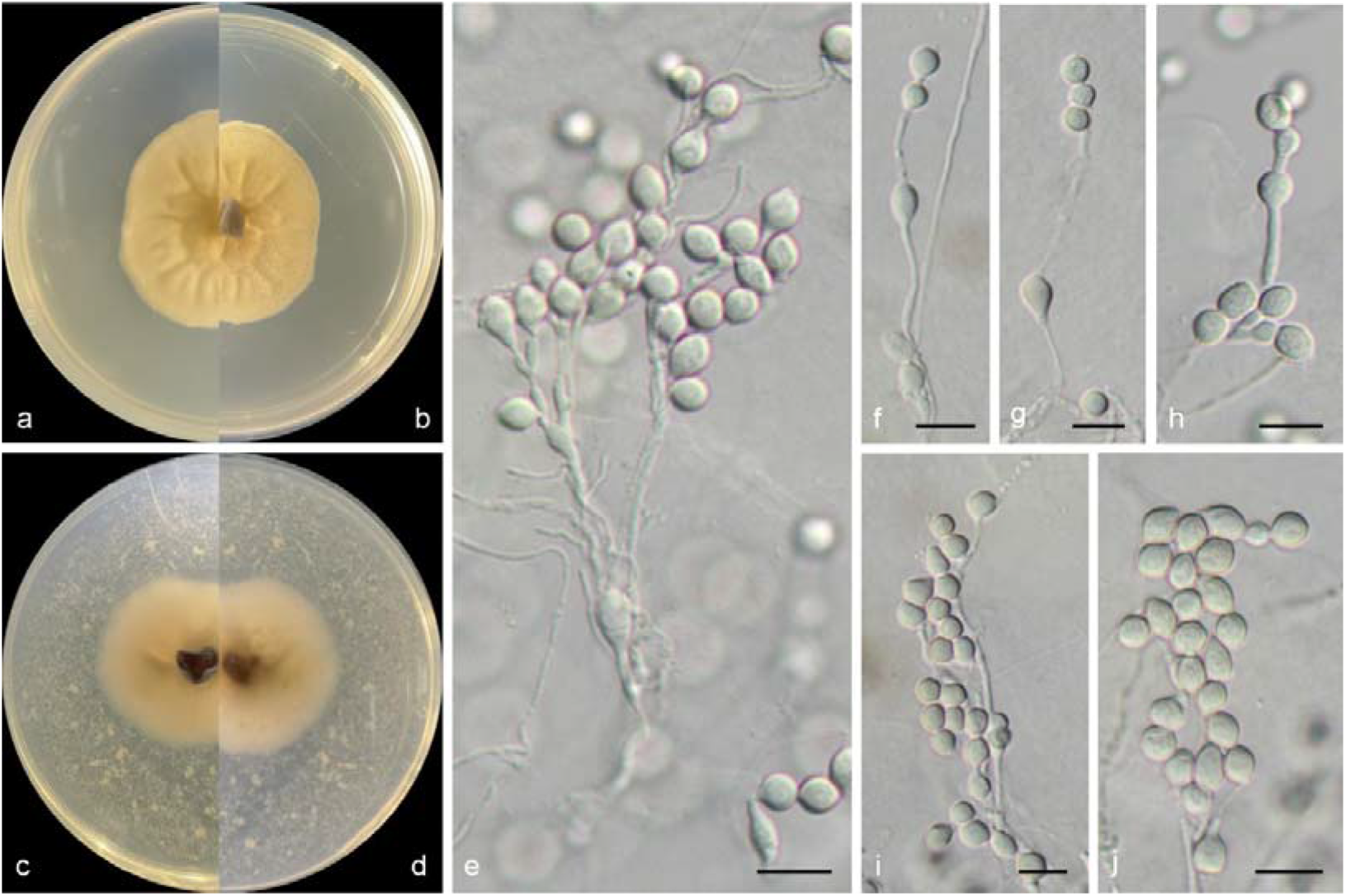
morphological photograph of *Chrysosporium sphaerospermum* (ex-type OUCMB□_14_0926). a-d: colonies on PDA and CDA after 7d; e-h: Conidiophores and conidiogenous cells; i-j: Conidia. Bars: e-j = 10 μm.

*Etymology*: named after the shape of conidia of this species, near globose.

*Typus*: CHINA, Shandong Province, Dongying city (119°09′45.18″E 37°45′34.86″N), from sediment, Jun. 2014, M.M. Wang (holotype designated here, dried culture on SNA; culture ex-type CGMCCXXXX = WL02839).

Vegetative hyphae hyaline, septate, branched, smooth. Ascomata not observed. Peridial hyphae, septate, branched and anastomosed, smooth, thick-walled, hyaline, 2–3 µm diam. Arthroconidia abundant, intercalary, laterial or terminal, unicellular, hyaline; intercalary conidia near globose, rarely racquet, with truncated base, 5–8 × 4–5 µm; laterial or terminal conidia arised from aerial hyphae directly, near globose to pyriform, with truncated base, 5–8 × 4–5 µm.

Culture characteristics — Colonies on PDA 30–35 mm diam in 7 d at 25°C, flat, fold deformation, aerial hyphae absent, pale brown near the centre, with white regular margin; reverse pale brown near the centre, white near the margin. Colonies on CDA 25–30 mm diam in 7 d at 25°C, flat, pale brown in the centre, with white regular margin; reverse pale brown near the centre, white near the margin.

Other examined isolates: CHINA, Fujian Province, Putian city (119°07′19.94″E 25°24′32.70″N), from sediment, May. 2014, X.M. Bian and W. Li (WL02687); Guangdong Province, Shantou city (116°45′29.03″E 23°18′37.00″N), from sediment, May. 2014, X.M. Bian and W. Li (WL02759).

## Discussion

In this study, three new species isolated from coastal sediments of China were described and illustrated, namely *Aphanoascus sedimenticola, Chrysosporium microsporum* and *C. sphaerospermum*. Phylogenetically, *A. sedimenticola* formed a single clade in the genus *Aphanoascus*, closest related to *A. jingzhouensis, A. keratinophilus* and *A. reticulisporus* (Fig. 1). Morphologically, *A. sedimenticola* could be distinguished in the shape and size of conidia (racquet or clavate with truncated base, 5–7 × 2–4 µm in *A. sedimenticola* vs. tubby to oblong-clavate, 4.3–32.4 × 2.2–7.6 µm in *A. jingzhouensis*, pyriform, 8.5–13 × 5.5–9 µm in *A. keratinophilus*, and conidia not observed in *A. reticulisporus*) (Routien 1967; Cano and Guarro 1990; Zhang et al. 2017). Although *C. microsporum* and *C. sphaerospermum* were close in terms of phylogenetical relationship (Fig. 1), they could be separated by the morphologic characteristics (e.g., shape and size) of conidia. For *C. microsporum*, it has cylindrical, ellipsoidal, pyriform or clavate conidia with the size of 3–5 × 2.5–3 µm, and for the latter, *C. sphaerospermum* is identified by near globose to pyriform conidia, rarely racquet, with the size of 5–8 × 4–5 µm.

